# EndD mediates butyrate-dependent toxin release in *Clostridioides difficile*

**DOI:** 10.64898/2026.06.22.733756

**Authors:** Horia A. Dobrila, Henrieta Licha, Andrew J. Hryckowian

## Abstract

*Clostridioides difficile* is an urgent threat to human health. Current treatments for *C. difficile* infections (CDIs) are antibiotics and microbiome restoration therapy (MRT) for recurrent cases. However, antibiotics contribute to antibiotic resistance and recurrent CDIs and the long-term sustainability and accessibility of MRTs remains to be determined. Since a dysbiotic gut microbiome is the primary risk factor for CDI, a better understanding of the interactions between *C. difficile*, the microbiome, and the host will aid development of treatments with improved precision. Emerging evidence supports that butyrate, a prominent end product of gut microbiome metabolism, is a key determinant of *C. difficile* pathogenesis. Notably, *C. difficile* releases more of its toxins TcdA and TcdB in butyrate-rich environments. Here, we demonstrate that butyrate-dependent toxin release is not driven by two previously characterized modes of toxin release (e.g., TcdE-dependent secretion or Cwp19-dependent autolysis). Instead, butyrate enhances the expression of a broadly conserved endolysin (EndD), which is responsible for butyrate-dependent toxin release. We additionally demonstrate that *endD*-dependent toxin release does not universally occur under all growth conditions and that its expression is dependent on the late-stage sporulation sigma factor SigK. Overall, our findings provide deeper insight into butyrate-dependent effects on *C. difficile* pathogenesis and set the stage for future work to better understand the molecular and genetic underpinnings of *endD* regulation.

## Introduction

The Centers for Disease Control and Prevention classifies *C. difficile* as an urgent threat to the nation’s health, resulting in 450,000 infections, 15,000 deaths, and 1 billion dollars in healthcare costs per year in the United States^1,2^. Most *C. difficile* infections (CDIs) occur in healthcare settings, where CDI is the most common cause of infectious diarrhea^3^. Risk factors for CDI include antibiotics, proton pump inhibitors, impaired immune function, advanced age, and diet, all of which are associated with dysbiotic gastrointestinal (GI) microbiomes^4^. While most infections are associated with antibiotic treatment, 22% of individuals with community acquired CDI have no recent history of antibiotic use. Additionally, factors affecting persistent and recurrent CDIs remain poorly defined^5,6^. Despite the morbidity and mortality caused by CDI, up to 15% of healthy adults are asymptomatic carriers of toxigenic *C. difficile*^7^, highlighting the gaps in our understanding.

*C. difficile* produces toxins and spores that are critical for its pathogenic lifecycle. TcdA and TcdB are protein toxins which it releases into the gut lumen. These toxins are internalized by colonocytes where they inactivate Rho GTPases, leading to the disruption of the actin cytoskeleton and tight junctions causing cell lysis, inflammation, and a weakened gut epithelial barrier characteristic of CDI^8–11^. The toxins are encoded within a 19.6 kb Pathogenicity Locus (PaLoc)^12^, also containing three accessory genes involved in toxin regulation (the sigma factor, *tcdR*^13^, and anti-sigma factor, *tcdC*^14^) and release (*tcdE*^15,16^). *tcdE* encodes a phage-like holin within the PaLoc, was originally hypothesized to be solely responsible for toxin release as a *tcdE* mutant was deficient in toxin release^15^ and exhibited reduced infection severity in a mouse model^16^. *C. difficile* can also release its toxins via autolysis^17^. The addition of glucose to growth media induces expression of *cwp19,* encoding a cell wall-degrading enzyme, and led to toxin release via autolysis^18^. Emerging evidence suggests that *C. difficile* toxin-mediated inflammation benefits *C. difficile* by allowing it to acquire host nutrients and outcompete inflammation-sensitive gut microbes^19–21^. *C. difficile* relies on spores to transmit between hosts. Regulation of sporulation is regulated by four sigma factors that coordinate the *C. difficile* cell’s asymmetrical division, forespore engulfment, assembly of the cortex/spore coat, and ultimate cell lysis of the mother cell to release the environmentally resistant spore^22–24^. Spores are resistant to heat, chemicals, desiccation, oxygen, and ethanol-based sanitizers^25^ allowing *C. difficile* to exist outside the GI tract and transmit to new hosts, where they germinate to form vegetative cells^26^.

Short chain fatty acids (SCFAs), in particular acetate, propionate, and butyrate, are metabolic byproducts of microbial degradation of dietary fiber and are the most concentrated metabolites in the distal gut, reaching concentrations >100 mM in the colon^27–29^. Gastrointestinal SCFA levels are influenced by microbiome composition, diet, antibiotics, and inflammation^30^. Of the SCFAs produced by the gut microbiome, butyrate stands out as an impactful metabolite as it is the main energy source for gut colonocytes^31,32^, promotes gut barrier integrity through several mechanisms^33–35^, and has direct immunomodulatory effects^36,37^. Animal and human data also show that high butyrate characterizes a gut that is non-permissive to CDI^38–41^. *In vitro*, butyrate suppresses *C. difficile* growth, increases sporulation, and leads to elevated *C. difficile* toxins in culture supernatants^35,38–40,42–45^. Together, these data suggest that *C. difficile* senses butyrate as a marker of a competitive GI environment and modulates its virulence to maintain a dysbiosis-associated niche or transmit to new hosts^46,47^.

While butyrate’s impact on *C. difficile* toxins is known, the genetic underpinnings of this phenotype are poorly characterized. Here, we confirm that butyrate has minimal impact on toxin transcripts, suggesting a difference in release, but not via TcdE nor Cwp19. Instead, we demonstrate butyrate-dependent expression of an endolysin, *endD*, is responsible for this phenotype and that the late-stage sporulation sigma factor, SigK, drives this expression.

## Results

### Exogenous butyrate does not alter toxin transcript abundance

Our previous work demonstrated via toxin-specific ELISAs and cell rounding assays that while exogenous butyrate elevates extracellular *C. difficile* toxin accumulation, it induces only minimal changes to transcripts within the Pathogencity Locus (PaLoc)^38,42^ To validate these findings, we performed reverse transcriptase quantitative PCR (RT-qPCR) on RNA extracted from cultures of *C. difficile* 630 grown mRCM with and without butyrate (**Figure S1**). These data confirm that across all three growth phases tested (mid-log, early stationary, and late stationary), there is no statistically significant increase in *tcdA* or *tcdB* transcripts. While many classical *C. difficile* toxin regulatory pathways operate at the transcriptional level^48^, our results demonstrate that butyrate alters extracellular toxin accumulation post-transcriptionally or by directly stimulating toxin release.

### Butyrate-mediated toxin release is not driven by TcdE or Cwp19

To determine if butyrate stimulates toxin accumulation through established pathways, we evaluated the roles of TcdE-mediated secretion and Cwp19-mediated autolysis. Clean, single-deletion mutants of *tcdE* and *cwp19* were generated in the *C. difficile* 630 strain background using a CRISPR-based mutagenesis system^49^. Because neither mutant exhibited baseline growth defects relative to the parental strain (**Figure S2AB**), we quantified extracellular toxin levels using cell rounding assays (**Figure 1A-C**). While both TcdE and Cwp19 are clearly important for efficient baseline toxin release (**Figure S3**), neither mutant abolished the butyrate-dependent toxin release phenotype (**Figure 1A-C, F**). Consistent with these findings, a double mutant (Δ*tcdE/*Δ*cwp19*) also retained its responsiveness to butyrate (**Figure S4**), confirming that butyrate drives toxin release via a non-classical mechanism.

**Figure 1.**
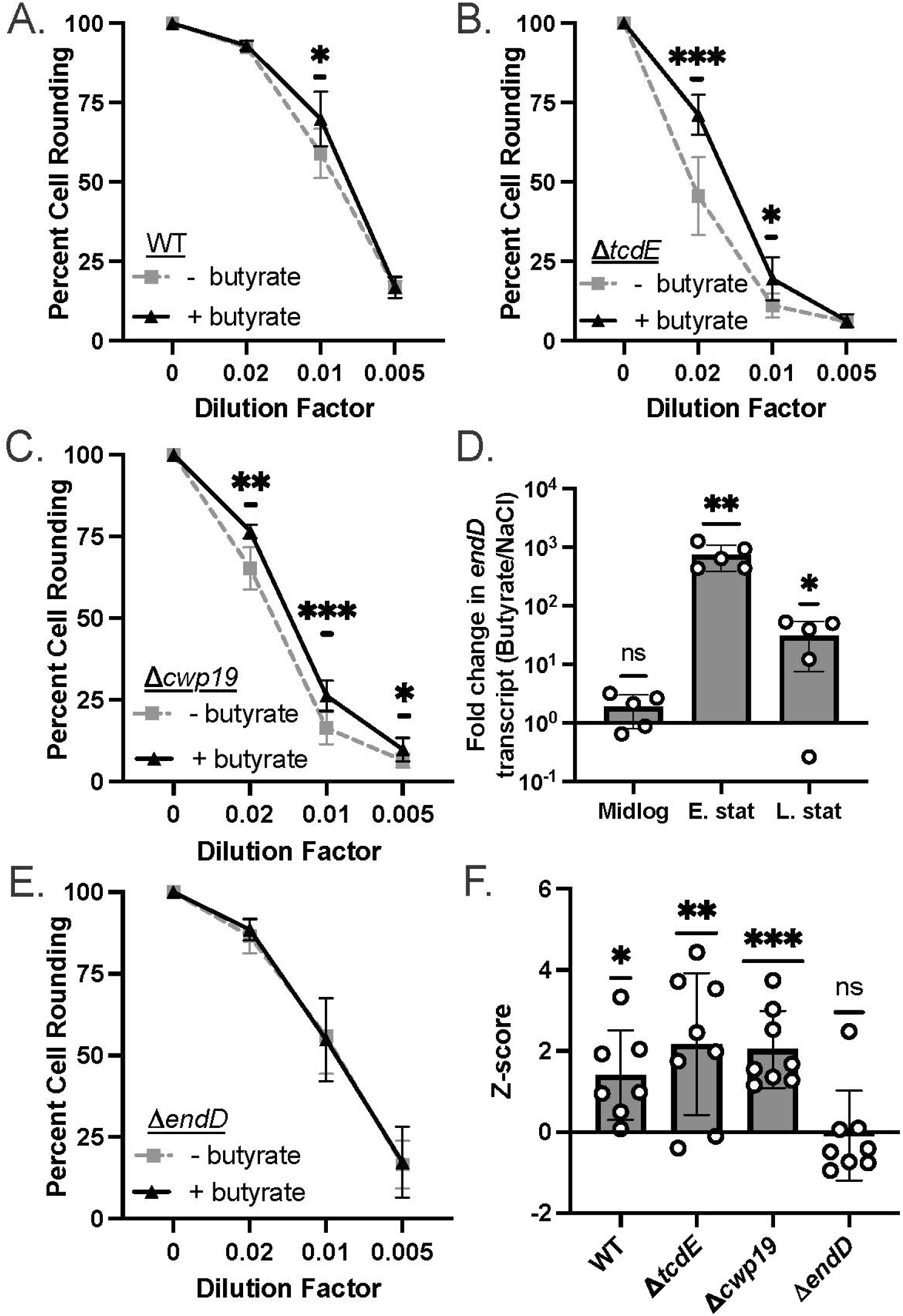
EndD is required for butyrate-dependent toxin release. Culture supernatants of *C. difficile* strains **(A)** 630 (WT) and isogenic **(B)** Δ*tcdE,* **(C)** Δ*cwp19,* and **(E)** Δ*endD* mutants grown in mRCM supplemented with either 50mM sodium chloride (- butyrate) or 50mM sodium butyrate (+ butyrate) were applied to HFFs to assay toxin abundance. Data points represent the mean percent total cell rounding at each supernatant dilution and error bars represent the standard deviation across n = 7 biological replicates for 630 and n = 8 biological replicates for the mutants. Statistical significance was determined at each dilution factor via Mann-Whitney test. **(D)** RNA was extracted from cultures of *C. difficile* strains 630 grown in mRCM supplemented with either 50mM sodium chloride or 50mM sodium butyrate at midlog, early stationary, and late stationary growth phases. Primers for *endD* were used to quantify transcript abundance compared to a housekeeping gene, *rpoC*, via RT-qPCR using the ΔΔCT method. Individual data points represent fold change of *endD* transcript abundance in butyrate supplemented cultures over control cultures, bars represent the mean value, and error bars represent the standard deviation across n = 5 biological replicates. Statistical significance was determined via one sample t-test with a hypothetical mean of 1. **(F)** Z-scores were calculated for butyrate-treated cultures relative to matched control cultures from the 1:100 supernatant dilutions from panels (A-C and E). Individual data points represent z-scores, bars represent the mean of individual data points, and error bars represent the standard deviation of data points. Statistical significance was determined via one sample t-test with a hypothetical mean of 0. *\*P* ≤ 0.05, *\*\*P* ≤ 0.01, *** *P* ≤ 0.001

### The endolysin EndD mediates butyrate-dependent toxin release

To identify candidate genes responsible for butyrate-dependent toxin release, we mined our previously published RNA-seq dataset for highly upregulated peptidoglycan-degrading enzymes^42^. From this, we identified CD630_21840, which encodes a predicted N-acetylmuramoyl-L-alanine amidase. This endolysin has 100% amino acid identity with a homolog in the hypervirulent M7404 strain (M7404_02200) that was recently shown to mediate toxin release in a non-lytic manner and is broadly conserved with >95% amino acid identity among diverse *C. difficile* clinical isolates^50^. To maintain consistency with established nomenclature of endolysins associated with large Clostridial toxin release in related species (e.g., *endS* in *P. sordelii* and *endP* in *C. perfringens*^50^), we designated this gene as *endD*. To validate our previous RNA-seq data^42^, we performed RT-qPCR on RNA extracted from *C. difficile* 630 grown in mRCM with and without butyrate. At both early and late stationary phase growth, *endD* transcripts were strongly upregulated in the presence of butyrate (**Figure 1D**).

Next, we constructed a clean deletion mutant (Δ*endD*) in the *C. difficile* 630 strain background. The mutant exhibited no baseline growth defects compared to wildtype (**Figure S2C**). In cell rounding assays, although the Δ*endD* strain showed a significant defect in butyrate-independent toxin release (**Figure S3**), it was small in comparison to defects in butyrate-independent toxin release observed in the Δ*tcdE* and Δ*cwp19* mutants (**Figure 1A-C, E, and Figure S3**). However, there was complete loss of the butyrate-dependent toxin release (**Figure 1EF**). To confirm that this defect was specific to the loss of *endD*, we complemented the mutant with a plasmid containing *endD* under the control of its native promoter. This plasmid-based complementation restored butyrate-dependent *endD* transcription during both early and late stationary time points (**Figure 2A**) and restored the butyrate-dependent toxin release phenotype (**Figure 2BC)**. These loss of function and complementation data demonstrate that EndD is required for butyrate-dependent toxin release in *C. difficile*.

**Figure 2.**
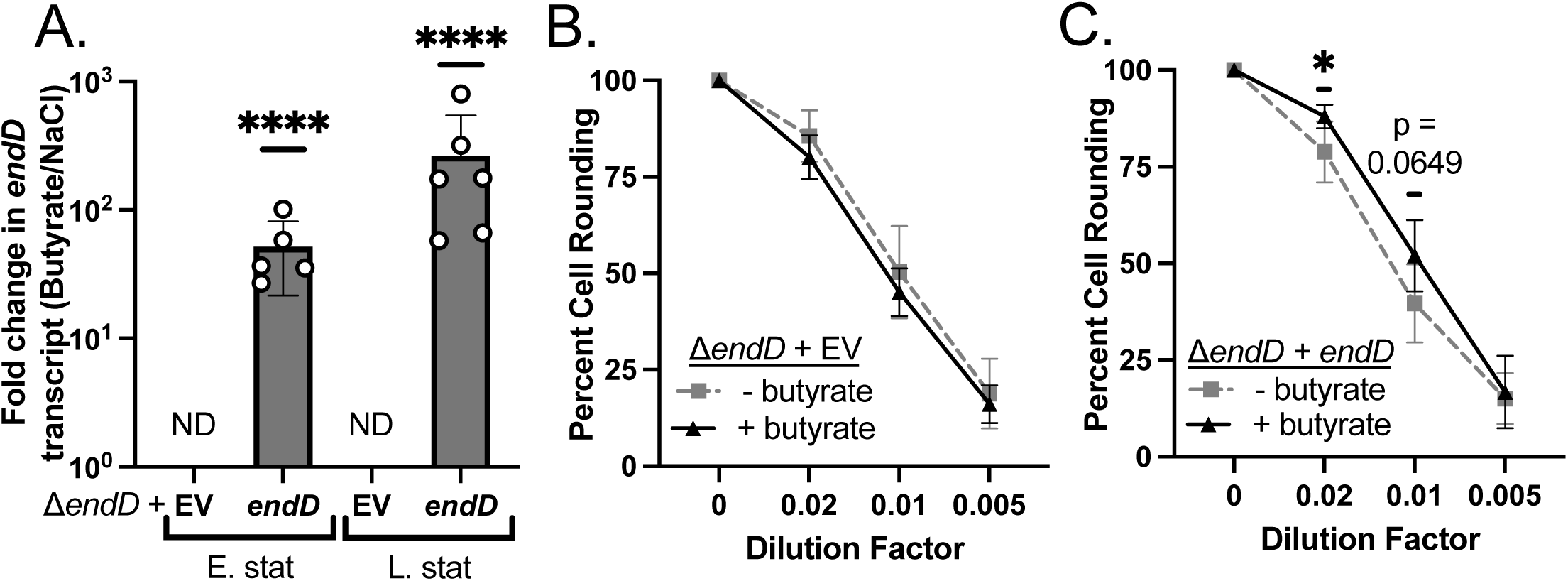
Genetic complementation of *endD*. **(A)** RNA was extracted from *C. difficile* 630 Δ*endD* containing either an empty vector (EV) or *endD* complementation (+*endD*) plasmid grown in mRCM + 10ug/mL thiamphenicol supplemented with either 50mM sodium chloride or 50mM sodium butyrate and sampled at early stationary and late stationary growth phases and RT-qPCR was performed using *endD*-specific primers to quantify transcript abundance compared to a housekeeping gene (*rpoC*) using the *ΔΔ*CT method. Individual data points represent fold change of *endD* transcript abundance in butyrate supplemented cultures over control cultures, bars represent the mean value, and error bars represent the standard deviation across n = 5 biological replicates. Statistical significance was determined via one sample t-test with a hypothetical mean of 1. Culture supernatants of *C. difficile ΔendD* containing either **(B)** EV or **(C)** *endD* complementation plasmid (*+endD*) were grown in mRCM + 10ug/mL thiamphenicol supplemented with either 50mM sodium chloride or 50mM sodium butyrate. Culture supernatants were applied to HFFs to assay toxin abundance. Individual data points represent mean percent total cell rounding and error bars represent the standard deviation across n = 6 biological replicates. Statistical significance was determined by Mann-Whitney test. \**P*≤ 0.05, *\*\*P* ≤ 0.01 and *\*\*\*\*P* ≤ 0.0001

### Butyrate-dependent toxin elevation via *endD* is nutrient-dependent

To determine the generalizability of butyrate-induced *endD* expression and its relationship to the to the toxin release phenotype, we evaluated these responses in a glucose-free basal defined medium (BDM), which was previously used to define butyrate-dependent growth defects in *C. difficile*^42^. We observed the butyrate-dependent toxin release phenotype in mRCM but not in BDM (**Figure 3AB**). Consistent with this loss of butyrate-dependent toxin release in BDM, we observed no butyrate-dependent elevation of *endD* transcripts at any of the time points tested (mid log, early stationary, late stationary) (**Figure 3C**). In line with the growing body of literature highlighting the importance of metabolic & environmental context in *C. difficile* toxin production and release mechanisms^48^, our results suggest that metabolic and environmental signals further impact the regulation butyrate-dependent of *endD* expression and toxin release.

**Figure 3.**
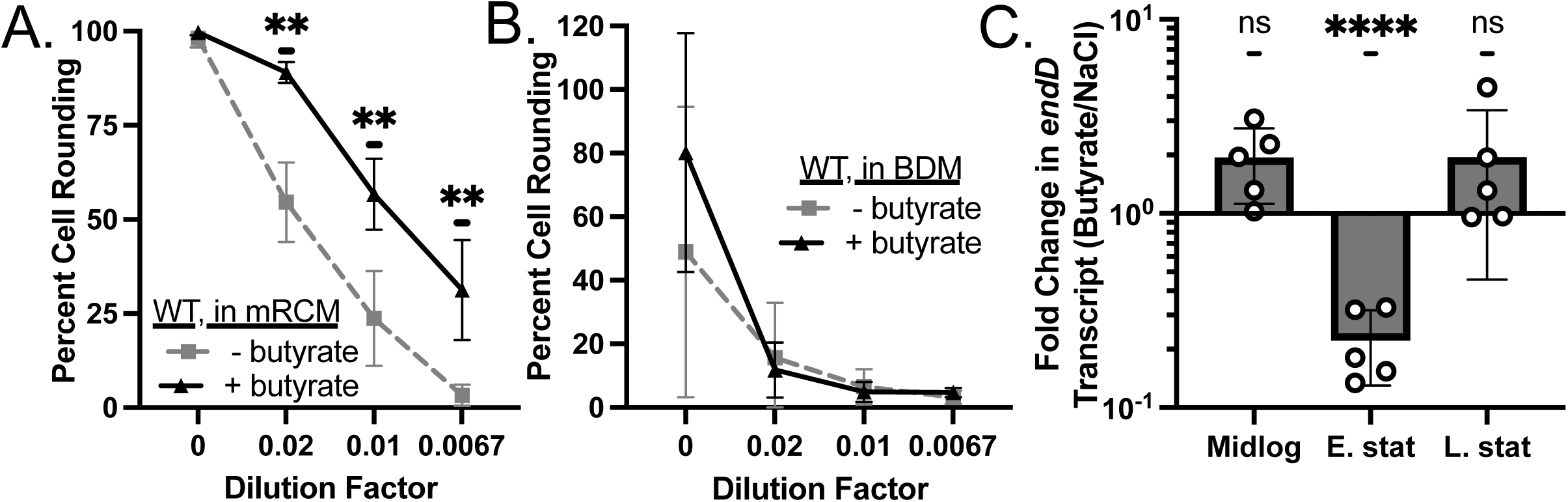
EndD-dependent toxin release does not occur in all growth media. Culture supernatants (48 hours post inoculation) of *C. difficile* 630 grown in **(A)** mRCM or **(B)** BDM supplemented with either 50mM sodium chloride or 50mM sodium butyrate were applied to human foreskin fibroblasts (HFFs) to assay toxin abundance via quantification of rounded cells. Individual data points represent percent total cell rounding, bar columns represent the mean value, and error bars represent the standard deviation across n = 6 biological replicates (for panel A) and n = 8 biological replicates (for panel B). Statistical significance was determined by Mann-Whitney test. **(C)** RNA was extracted from cultures of *C. difficile* 630 grown in BDM supplemented with either 50mM sodium chloride or 50mM sodium butyrate at midlog, early stationary, and late stationary phases of growth and used for RT-qPCR. Specific primers for *endD* were used to quantify transcript abundance compared to a housekeeping gene, *rpoC*, using the ΔΔCT method. Individual data points represent fold change of *endD* transcript abundance in butyrate supplemented cultures over control cultures, bars represent the mean value, and error bars represent the standard deviation across n = 5 biological replicates. Statistical significance was determined by one sample t-test with a hypothetical mean of 1. \**P*≤ 0.05, *\*\*P* ≤ 0.01 and *\*\*\*\*P* ≤ 0.0001.

### The sporulation sigma factor SigK regulates butyrate-dependent *endD* expression

While *endD’s* importance for toxin release is a recent discovery^50^, this gene was previously linked to *C. difficile* sporulation. Prior transcriptomic characterization of the sporulation sigma factors regulons noted reduced *endD* transcript abundance in *sigK*, *sigE*, and *sigF* mutants compared to wildtype^51,52^. Additionally, a consensus SigK promoter sequence is located upstream of the *endD* coding region^51–53^. To determine whether butyrate-dependent transcription of *endD* is mediated by SigK, we leveraged a *sigK* gene disruption mutant generated via Targetron^24^. The *sigK* mutant and its parental strain (*C. difficile* 630 Δ*erm)* strain were grown in mRCM with and without butyrate. Total RNA was harvested at early stationary and late stationary phases of growth for RT-qPCR analysis. We observed complete loss of butyrate-dependent *endD* expression in the *sigK* mutant at both time points (**Figure 4**), indicating that the mother-cell sigma factor SigK is strictly required for butyrate-dependent *endD* transcription.

**Figure 4.**
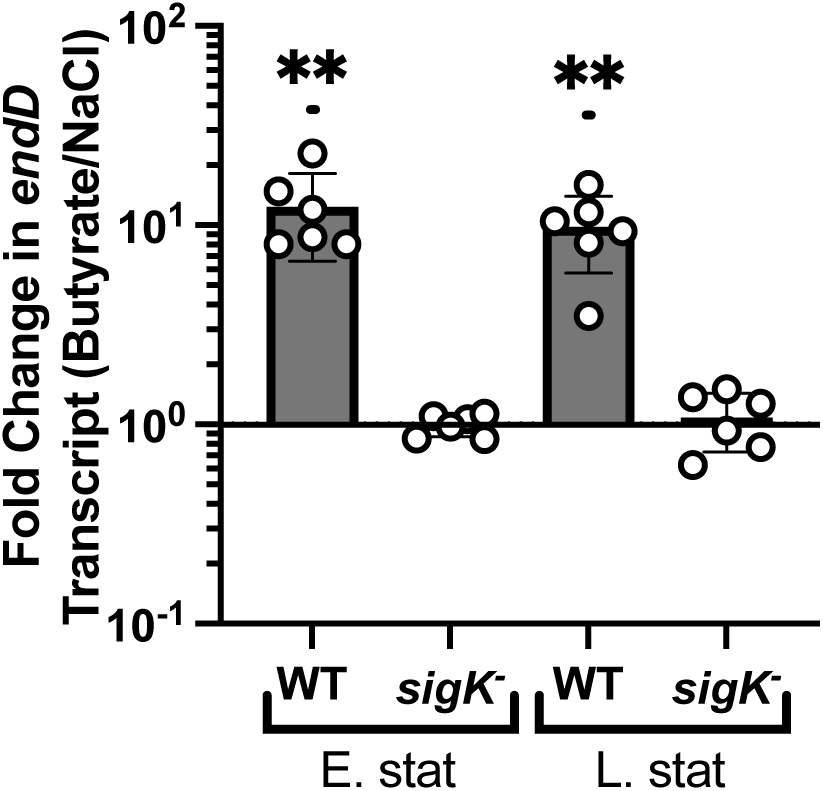
Transcriptional regulation of *endD* by SigK during butyrate exposure. RNA was extracted from cultures of *C. difficile* 630 Δ*erm* (WT) or an isogenic *sigK*- mutant grown in mRCM supplemented with either 50mM sodium chloride or 50mM sodium butyrate at early stationary and late stationary growth phases. Specific primers for *endD* were used to quantify transcript abundance compared to a housekeeping gene, *rpoC*, via RT-qPCR using the ΔΔCT method. Individual data points represent fold change of *endD* transcript abundance in butyrate supplemented cultures over control cultures, bars represent the mean value, and error bars represent the standard deviation across n = 6 biological replicates. *\*\*P* ≤ 0.01 via Mann-Whitney test.

## Discussion

Toxin accumulation is a well-established consequence of butyrate exposure in *C. difficile*^47^ but the molecular and genetic underpinnings of this phenotype have not been determined. In this work, we resolve a major piece of this regulatory puzzle. We demonstrate that while both *tcdE* and *cwp19* are important genes for general toxin release, neither is directly involved in the butyrate-dependent toxin release phenotype (**Figure 1**). Instead, we identify the endolysin EndD as an essential player in dictating this response (**Figures 1 and 2**). EndD broadly conserved with high sequence identity (>95% at the amino acid level) among *C. difficile* isolates^50^, highlighting the importance of its function in butyrate-dependent toxin release (**Figure 1E**) and to a lesser extent, butyrate-independent toxin release (**Figure S3;** which was also observed in *C. difficile* M7404^50^). In addition, it was previously hypothesized that EndD crosses the bacterial membrane via TcdE^50^, similar to holin-endolysin pairs which facilitate bacteriophage egress^54^. However, our data challenge this model. Because a *tcdE* deletion mutant still exhibits butyrate-dependent toxin release (**Figure 1B**), EndD may be delivered to the bacterial cell wall by an alternative transmembrane exporter. While further work is needed to understand the potential differences in butyrate-responsive toxin release between strains and elucidate the how EndD is exported, we have identified an endolysin that directly couples a prominent gut microbial metabolite to elevated *C. difficile* virulence.

A key finding of this study is that the EndD-mediated toxin release phenotype is dictated by the surrounding nutritional landscape. While butyrate drives toxin release in several growth media including mRCM, BHI, and CDMM^35,38,42,44,47^, this phenotype and butyrate-dependent *endD* transcription are abolished in BDM (**Figure 3**). Similarly, our previous work demonstrated that *C. difficile*’s characteristic butyrate-dependent growth defect is absent in BDM but that this growth defect worsens as additional nutrients are added to the medium^42^. Ongoing work in our laboratory seeks to define the co-occurrence versus divergence of the butyrate-dependent growth defect and toxin release phenotypes across controlled media conditions. As such, we expect to identify specific metabolic inputs that can decouple these phenotypes. Another possibility is that common regulatory circuitry underlies both phenotypes and as such, they are inextricably linked, and this ongoing work will identify these regulatory elements. Regardless, this continued investigation will provide insights into the molecular targets needed to selectively leverage butyrate to simultaneously minimize *C. difficile* levels and virulence in the GI tract.

Building off of existing literature that identified a putative SigK promoter site upstream of *endD*^53^, we demonstrate that butyrate- specific transcription of *endD* is mediated via the late-stage sporulation sigma factor, SigK (**Figure 4**). This highlights previously unappreciated intricacies of regulation between *C. difficile’s* toxin release and sporulation response phenotypes. Further study into the mechanistic underpinnings of SigK’s control over *endD*-mediated toxin release is essential to unraveling co-occurrence of this and the butyrate-dependent sporulation phenotype. Because *endD* is in the SigK regulon, EndD was previously suggested to facilitate mother cell lysis during sporulation^51,52^. Our results do not exclude this possibility, instead they suggest that EndD may have dual functionality. For example, despite the “division of labor” between sporulation and toxin gene expression in individual *C. difficile* cells within populations^55^, there may be additional regulatory checkpoints or cellular contexts that influence the function of EndD within individual cells. Regardless, this work begins to reframe SigK as a pleiotropic regulator of virulence and transmission in *C. difficile*.

## Materials & Methods

### Bacterial strains and culture conditions

All bacterial growth media were pre-reduced for a minimum of 24 hours in an anaerobic chamber prior to use in experiments, and all bacterial growth was performed under anaerobic conditions in an anaerobic chamber (Coy).

*C. difficile* strains (**Table S1**) were maintained as freezer stocks in 25% glycerol under anaerobic conditions in septum-topped vials at -80°C. *C. difficile* strains were routinely cultured anaerobically at 37°C on CDMN agar (*Clostridioides difficile* agar with moxalactam and norfloxacin), composed of *C. difficile* agar base (Oxoid) supplemented with 7% defibrinated horse blood (HemoStat Laboratories), 32 mg/L moxalactam (Santa Cruz Biotechnology), and 12 mg/L norfloxacin (Sigma-Aldrich). After 16-24 hours of growth on agar plates under anaerobic conditions, a single colony was picked into 5 mL of modified reinforced clostridial medium (mRCM, an undefined rich medium^42^) or basal defined medium (BDM, a defined medium lacking glucose and containing amino acids designed to support *C. difficile* growth^56^) and grown anaerobically at 37°C for 16-24 hrs. These overnight cultures from single colonies were used to subculture into necessary medium to be used in experiments.

For *in vitro* experiments examining bacterial growth, subcultures were prepared at 1:200 dilutions in mRCM or BDM supplemented with 50mM sodium butyrate or 50mM sodium chloride. After addition of these supplements and prior to pre-reduction of the media in an anaerobic chamber, the pH of the media was adjusted to pH 6.5 and filter-sterilized using 0.2 µm PVDF filters (Thermo Fisher Scientific). Growth curve experiments were done in sterile polystyrene 96-well tissue culture plates with low evaporation lids (Falcon). Cultures were grown anaerobically in a BioTek Epoch2 plate reader. At 30-min intervals, the plate was shaken on the “slow” setting for 1 min and the OD600 of the cultures was recorded using Gen5 software (version 1.11.5).

### Genetic modification of *C. difficile*

See **Tables S1-S3** for information on strains, plasmids, and oligonucleotides. Clean gene deletions were created in *C. difficile* strain 630 using a single plasmid, xylose-inducible CRISPR-*cas9* system as previously described^49^. Briefly, homology regions (300-500bps) corresponding to the upstream and downstream regions of the gene of interest (including start and stop codons of the gene) were PCR-amplified and assembled into pIA123. Three to five sgRNAs were designed per gene of interest using Benchling’s built-in CRISPR guide design tool using the following parameters: single guide; guide length = 20; genome = ASM920v1 (*Peptoclostridium difficile*); PAM = NGG (SpCas9, 3’ side). Guides were picked based on highest “on-target” and “off-target” scores from this tool. The selected guides were then assembled into the homology region-containing plasmid, resulting in 3-5 plasmids containing the same homology region and different sgRNAs. These plasmids were sequence confirmed, transformed into a conjugation proficient strain of *E. coli* (HB101/pRK24) and conjugated into *C. difficile*. Mutagenesis was induced with 1% xylose, after which candidate colonies were cured of the plasmid. Sanger sequencing was done to confirm loss of the gene of interest.

Complementation of *endD in trans* was performed using a modified version of pAP114^57^, where the xylose repressor and promoter sequence were removed and *endD* plus 300bps upstream of *endD* were cloned in between the SacI and BamHI restriction enzyme sites, to ensure expression occurs under native promoter control rather than xylose-inducible control. This construct and an empty vector control were conjugated into the *ΔendD* background and maintained by supplementing all necessary media with 10 *μ*g/mL of thiamphenicol.

### *C. difficile* toxin quantification

*C. difficile* toxins were quantified in 48 hour culture supernatants using a cell rounding assay as previously discussed^42^. Specifically, two days before cell treatment, overnight cultures of *C. difficile* grown in mRCM or BDM were back-diluted 1:200 into mRCM containing 50mM sodium butyrate or 50mM sodium chloride (pH adjusted to pH 6.5). Cultures were grown anaerobically at 37°C for 48 hrs. One day before treatment, confluent human foreskin fibroblast (HFF) cells (ATCC SCRC-1041) grown in HFF medium (see below for HFF medium composition) were harvested and counted with a hemocytometer and seeded into a 96-well plate at 8,000 cells per well and incubated at 37C under 5% CO2 for 24 hrs.

On the day of experiments, *C. difficile* supernatants were centrifuged (3,000 x g for 10 minutes at room temperature) and filter sterilized. Butyrate was removed from culture supernatants by size exclusion filtration to minimize its protective effects against *C. difficile* toxins in eukaryotic cells^35^. Specifically, 5mL of filtered culture supernatant was transferred to Pierce 100kDa MWCO filters and was centrifuged at 3,000 x g for 5 minutes at room temperature. The filter dead volume (containing molecules > 100kDa, including *C. difficile* toxins) was then washed with 5mL of 1x PBS and was centrifuged again at 3,000 x g for 5 mins. Then the washed fraction containing C. difficile toxins was reconstituted in 5mL of HFF medium to dilute this fraction back to original concentration found in culture supernatants. Concentrations of the resulting toxin-containing HFF media were prepared at undiluted, 1:50, 1:100, and 1:150 or 1:200 dilutions. Phase contrast images were acquired using Sartorius Incucyte to confirm the health and morphology of the HFF cells prior to treatment. Then, the medium was removed from the adherent HFF cells and toxin containing media at the dilutions noted above were added. Cells were incubated with toxin containing media for 4 hours and imaged again. Rounded cells and unrounded cells were manually counted.

HFF medium contains the following components by volume: 88% high glucose DMEM (Thermo Scientific 11965126), 10% heat-inactivated fetal bovine serum, 1% penicillin/streptomycin (Thermo Scientific 15140122), and 1% Glutamax (L-alanyl-L-glutamine) (Thermo Scientific 35050061).

### Reverse Transcription – Quantitative PCR (RT-qPCR)

Cultures of *C. difficile* 630 were grown in pre-reduced mRCM or BDM and sampled at midlog phase growth (OD600 = 0.3-0.5 in mRCM or OD600 = 0.18-0.2 in BDM), early stationary phase growth (24 hours post inoculation), and late stationary phase growth (48 hours post inoculation). At each time point, bacterial culture was taken and immediately mixed with an equal volume of ice cold 1:1 acetone/ethanol mixture and placed at -20°C prior to preserve RNA. RNA extraction was performed as previously described^42^. Briefly, samples were centrifuged at 3,000 x g for 5 minutes at 4°C. The supernatant was removed, pellets were washed with 5mL of cold, nuclease-free PBS and were centrifuged at 3,000 x g for 5 minutes at 4°C. The supernatant was removed, and remaining pellets were resuspended in 1mL of TRIzol and processed using a TRIzol Plus RNA Purification Kit (Thermo) with on-column DNase treatment according to the manufacturer’s instructions. Purified RNA was frozen at -80°C.

RT-qPCR was performed on extracted RNA using the GoTaq 1-Step RT-qPCR kit (Promega) according to manufacturer’s instructions, with a final amount of 40 ng of RNA in a final volume of 10 µL. Each reaction was setup with primers amplifying the gene(s) of interest along with *rpoC* (as a housekeeping gene) and was run with three technical replicates. The RT-qPCR steps were performed on a QuantStudio 7 Flex (Applied Biosystems), and the threshold cycle (Ct) was determined using QuantStudio Real-Time PCR Software v1.7.2. The cycle run involved an initial reverse transcriptase activity step of 40°C for 15 mins followed by an inactivation step of 95°C for 10 minutes. The PCR cycles were 95°C for 10 seconds, 60°C for 30 seconds, and 72°C for 30 seconds for a total of 40 cycles, with a melt curve performed afterwards. Relative fold change was determined using the 2^-ΔΔCT^ method by comparing the Ct values in the presence and absence of butyrate within each strain.

### Statistical analysis

Statistical analysis was performed using GraphPad Prism 11.0.1. Details of specific analyses, including statistical tests used, are found in applicable figure legends. * = *P* ≤ 0.05, ** = *P* ≤ 0.01, *** = *P* ≤ 0.001, **** = *P* ≤ 0.0001.

## Acknowledgements

This material is based upon work supported by NIH R01 AI195616 (to A.J.H), the NSF GRFP (DGE-2137424 to H.A.D.), and startup funding from the University of Wisconsin-Madison (A.J.H.). Any opinions, findings, and conclusions or recommendations expressed in this material are those of the authors and do not necessarily reflect the views of the funders. A.J.H. is the Judy L. and Sal A. Troia Professor in Gastrointestinal Disease Research at the University of Wisconsin-Madison.

H.A.D. and H.L. performed experiments and analyzed the data. H.A.D. and A.J.H. prepared the display items. H.A.D. and A.J.H. wrote the paper. All authors edited the manuscript prior to submission.

